# GP-HTNLoc: A Graph Prototype Head-Tail Network-based Model for Multi-label Subcellular Localization Prediction of ncRNAs

**DOI:** 10.1101/2024.03.04.583439

**Authors:** Shuangkai Han, Lin Liu

## Abstract

Numerous research findings demonstrated that understanding the subcellular localization of non-coding RNAs (ncRNAs) is pivotal in elucidating their roles and regulatory mechanisms in cells. Despite the existence of over ten computational models dedicated to predicting the subcellular localization of ncRNAs, a majority of these models are designed solely for single-label prediction. In reality, ncRNAs often exhibit localization across multiple subcellular compartments. Furthermore, the existing multi-label localization prediction models are insufficient in addressing the challenges posed by the scarcity of training samples and class imbalance in ncRNA dataset. This study addresses the limitations of existing models by introducing a novel multi-label localization prediction model for ncRNAs, termed GP-HTNLoc. To alleviate class imbalance, the model adopts a separate training approach for head and tail class labels. In GP-HTNLoc, a pioneering graph prototype module is introduced for capturing potential association of ncRNA samples with labels. This module efficiently learns the graph structure and aggregates sample features. Notably, only few samples are required to obtain label prototypes containing rich information. These prototypes are then utilized to train a transfer learner, facilitating the transfer of meta-knowledge from the head class to the tail class. Experimental results demonstrate that GP-HTNLoc surpasses current state-of-the-art models across all datasets. Ablation study underscore the vital role played by the graph prototype module in enhancing the performance of GP-HTNLoc. The user-friendly online GP-HTNLoc web server can be accessed at https://56s8y85390.goho.co.

## 1 Introduction

Previous studies have identified a large number of non-coding RNAs (ncRNAs) in the mammalian genome, and while it is entirely possible that most of these ncRNAs are transcriptional noise or by-products of RNA processing, there is growing evidence that most of them are functional and provide a variety of regulatory activities in the cell [1].Recent research underscores the intimate association between ncRNAs and the onset and progression of specific diseases [2]. Particularly, most ncRNAs exhibit varying local concentrations, interacting partners, post-transcriptional modifications, and regulatory pathways in diverse subcellular locations. These differences significantly impact protein synthesis and cellular functions [3,4]. For instance, miR-122, a highly expressed miRNA in the liver, predominantly functions within the cytoplasm of hepatic cells. Its influence on metabolic activities in the liver arises from its binding to target gene mRNAs, thereby regulating their translation or stability [5]. Consequently, the investigation of the subcellular localization of ncRNAs emerges as a crucial avenue for unraveling the functional intricacies and regulatory mechanisms inherent in these molecules. This work provides substantial value for researchers in elucidating gene regulation, cellular functions, and the mechanisms underlying various diseases in organisms.

Although traditional RNA subcellular localization methods can ensure high localization accuracy, they are only suitable for small-scale studies because they are expensive and time-consuming [6]. Facing the current high-throughput needs, researchers are trying to find some computational methods to replace traditional methods for ncRNA subcellular localization. Supported by a rapidly growing database of ncRNA subcellular localization[7,8,9,10], computational model-based methods for subcellular localization of ncRNAs have emerged as a focal point of this research domain in recent years.

Non-coding RNAs (ncRNAs) are classified into different categories according to their length, function, and subcellular location, and the ncRNAs that have been widely studied by the biological community include long noncoding RNAs (lncRNAs), small nucleolar RNAs (snoRNAs) and microRNAs (miRNAs) [11]. For lncRNAs, Fan et al. devised lncLocPred, a logistic regression-based machine learning predictor dedicated to predicting the subcellular localization of lncRNAs [12]. Addressing the challenge of limited samples in lncRNA subcellular localization, Cai et al. presented GM-lncLoc, a meta-learning training model facilitating knowledge transfer through meta-parameters [13]. Regarding miRNAs, Yang et al. developed MiRGOFS, a functional similarity measure based on Gene Ontology (GO), for predicting miRNA subcellular localization and miRNA-disease associations [14]. Xiao et al. proposed miRLocator, a sequence-to-sequence learning model with an attention mechanism for identifying human miRNA subcellular localization [15]. Wang et al. applied the Hilbert-Schmidt Independence Criterion for Multi-Kernel Learning (MK-HSIC), combining multivariate information and selecting optimal combinations for subcellular localization in the Support Vector Machine [16]. On this basis, Zhou et al. introduced the MKGHKNN model, a multi-kernel graph regularization K local hyperplane distance nearest neighbor model [17]. Besides, Bai et al. presented the ncRNALocate-EL model, utilizing natural language processing to extract high-level features from ncRNA sequences. Overall, various machine learning models are currently blossoming in the field of ncRNA subcellular localization prediction [18].

Nonetheless, current ncRNA subcellular localization studies still face three major challenges: (1) Few-shot Challenge: Datasets containing reliable localization information for ncRNAs typically comprise only a few hundred sequences, insufficient for the effective training of machine learning models; (2) the class imbalance challenge: there are significant differences in the number of ncRNA samples involved in different subcellular locations, which makes it very difficult for the model to learn at locations where the number of samples is scarce; (3) Multi-labeling Challenge: Research indicates that an RNA sequence may exhibit multiple subcellular localizations [19], meaning that ncRNA subcellular localization is essentially a multi-label classification problem. However, the majority of existing studies focus on single-label localization for ncRNAs. Among the existing predictive models for ncRNA subcellular localization, several resampling techniques are employed in several enhanced machine learning-based localization prediction models to tackle class imbalance [20,21,22]. Although effective in achieving balance in the number of categories within the training data, these techniques come at the expense of losing the original data distribution features or introducing new noise. In addressing the multi-labeling challenge, a predominant approach employed by the majority of extant methodologies involves the utilization of the One-vs-Rest strategy [16,17,18]. However, the One-vs-Rest strategy overlooks the relationships between labels, often making it challenging for the model to learn label association information.

In multi-label classification problems in natural language processing (NLP), there is a significant class imbalance. A few of the labels (called head labels) are associated with a large number of documents, while the majority of the labels (called tail labels) are associated with a small number of documents, and the label frequency exhibits a distinct long-tailed distribution. This makes the tail class samples scarce and the model difficult to train [23,24,25]. To address this problem, Xiao et al. proposed the HTTN model, which employs a transfer learner that transforms class prototypes into classifier parameters, facilitating the transfer of meta-knowledge from data-rich head labels to data-poor tail labels [23]. The method of computing the average of multiple samples of the same class in HTTN to represent the prototype of the class is effective in the field of NLP where the amount of sample data is rich. However, for the problem of ncRNA subcellular localization prediction where both head and tail class samples are scarce, this method often fails to obtain a high-quality prototype representation, making it difficult to improve the subsequent classification accuracy. To address the challenge, this study proposes a novel model, GP-HTNLoc. The model employs the concept of transfer learning, leveraging meta-knowledge acquired from the data-rich head class to enhance the learning process in the data-scarce tail class. A key component of the model is the innovative graph prototype module. In this module, a Heterogeneous Graph is constructed using the association information between each ncRNA sample and its localization label. On this Heterogeneous Graph, diverse label-sample associations are gathered through Heterogeneous Graph Convolutional Networks (HGCN) [26] and the classical random walk algorithm MetaPath2Vec [27]. This process extracts the association information between various labels and samples from the intricate graph structure, allowing the model to learn embeddings for each label node. The graph prototype module requires only few samples to learn labeled prototypes containing rich graph structure information and potential label association information. These prototypes are then utilized to train the transfer learner, which maps the classification knowledge from header classes to tail classes, overcoming the class imbalance problem and constructing a comprehensive multi-label classifier.

To evaluate the performance of GP-HTNLoc, this study conducted a comparative analysis with state-of-the-art RNA subcellular multi-label localization models using publicly available benchmark datasets. Cross-validation results show that GP-HTNLoc exhibits significant performance advantages over existing models. To elucidate the impact of the graph prototype module in GP-HTNLoc, an ablation study of the component was conducted, and the results showed that the graph prototype module plays an important role in improving the performance of GP-HTNLoc.

## 2 Materials and Methods

### 2.1 Benchmark Dataset

In this study, we utilized multi-location non-coding RNA sequences derived from the RNALocate [28] database by Zhou and Wang [16,17] et al. as the foundational dataset for training and evaluating the model’s effectiveness. The dataset encompasses three non-coding RNA categories. These categories collectively span nine subcellular locations. Human-specific non-coding RNAs were exclusively extracted from the aforementioned three datasets, resulting in three independent datasets denoted as H_snoRNA, H_lncRNA, and H_miRNA. The distribution of sequence counts in each dataset is shown in Figure 1, where it can be seen that there is a significant class imbalance in each RNA subcellular localization dataset, with subcellular locations involving the most sequences accounting for nearly half of the total number of sequences, while subcellular locations involving the fewest sequences accounted for less than one-tenth of the total number of sequences.

**Figure 1:**
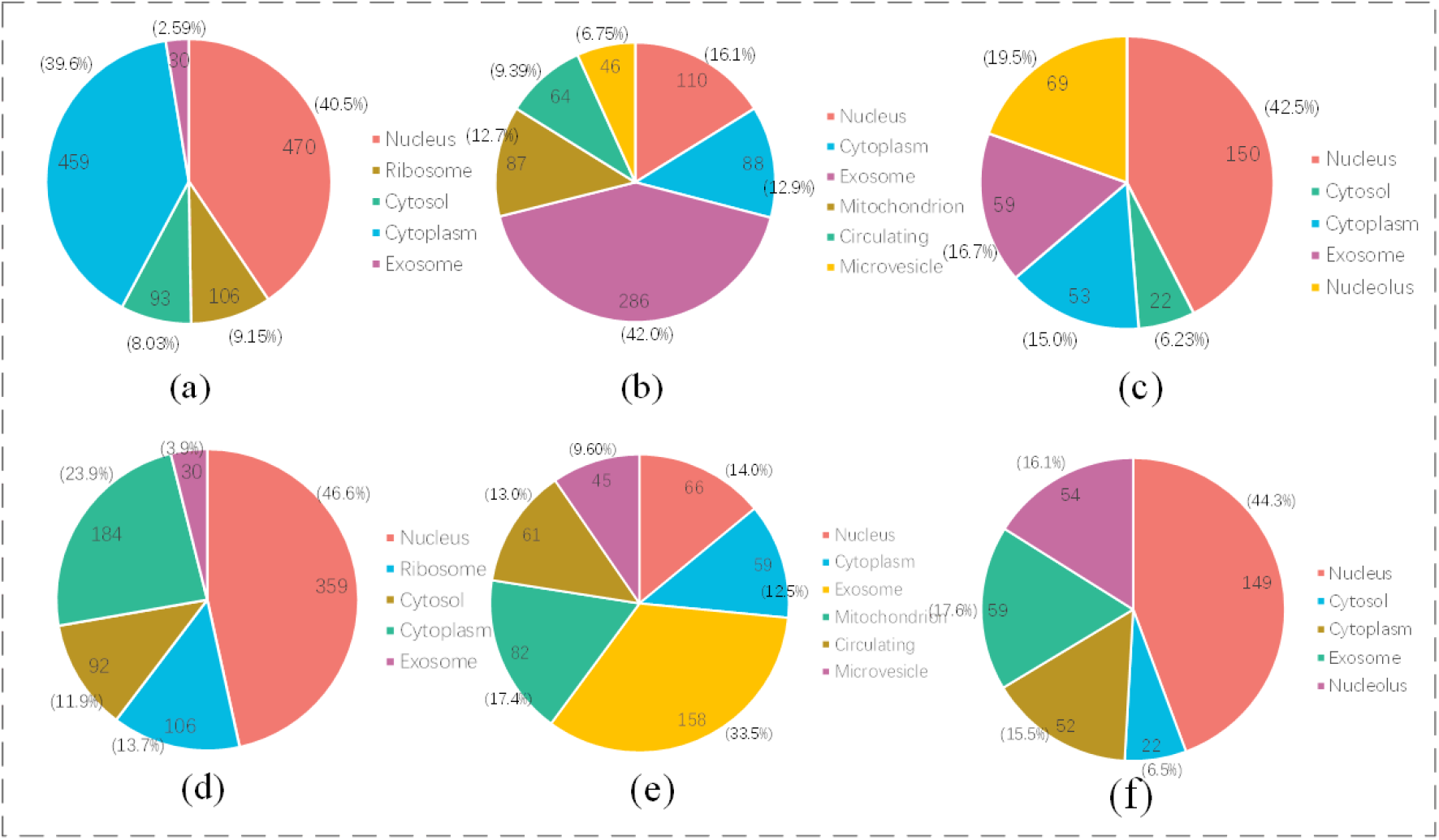
The distribution of subcellular on (a)LncRNA dataset; (b)miRNA dataset; (c)snoRNA dataset; (d)Human LncRNA dataset; (e)Human miRNA dataset; (f)Human snoRNA dataset.

### 2.2 Model Architecture

The overall architecture of the GP-HTNLoc model proposed in this study consists of three stages: (i) Imbalanced learning based on graph prototypes; (ii) Fine-tuning; and (iii) Prediction. In the first stage, the primary features of the sequence, denoted as *X*^*S*^, are first obtained by multiple sequence feature extraction methods. The set of *X*^*S*^ and its corresponding label dataset in the training set are referred to as the Base Data. The Base Data is then partitioned into *D*_*head*_ and *D*_*tail*_ based on head and tail labels, respectively.

For *D*_*head*_, on the one hand, the primary sequence features *X*^*S*^ is fed into a bidirectional long short-term memory (BiLSTM) network [33] to obtain advanced features represented as 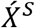. The head classifier parameters, denoted as *M*_*head*_, are subsequently trained using 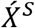. On the other hand, its primary features are directly fed into the proposed graph prototype module in this study. The graph prototype module utilizes the association information between ncRNA samples and labels to construct a heterogeneous graph *G*. On *G*, HGCN and MetaPath2Vec are used to learn the graph structure and aggregate sample features to obtain the embedding of labeled nodes, denoted as 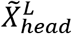. The transfer learner is then used to learn the mapping from 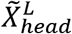 to *M*_*head*_.

For *D*_*tail*_, the primary features are input into the graph prototype module to obtain the prototype representation of tail class, denoted as 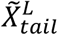. By inputting 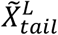 into the pre-trained Transfer Learner, the parameters for the tail classifier, denoted as *M*_*tail*_, are obtained. Finally, *M*_*head*_ and *M*_*tail*_ are concatenated to form the complete parameters *M* for ncRNA multi-label classification.

In the fine-tuning phase, *M* is lightly trained using tail class samples containing both head and tail labels, called Novel Data. In the prediction stage, after the new ncRNA sequences are acquired with deep sequence features, they are directly input into the fine-tuned *M* to obtain the prediction of subcellular multi-label localization of ncRNA sequences.

Figure 2 illustrates the overall workflow of GP-HTNLoc, using the imbalance learning, fine-tuning, and prediction on the LncRNA dataset as an example. In the whole model architecture, the graph prototype module plays a crucial role in training the transfer learner and the multi-label classifier. Subsequently, this paper describes the graph prototype module and overall workflow of GP HTNLoc in detail.

**Figure 2.**
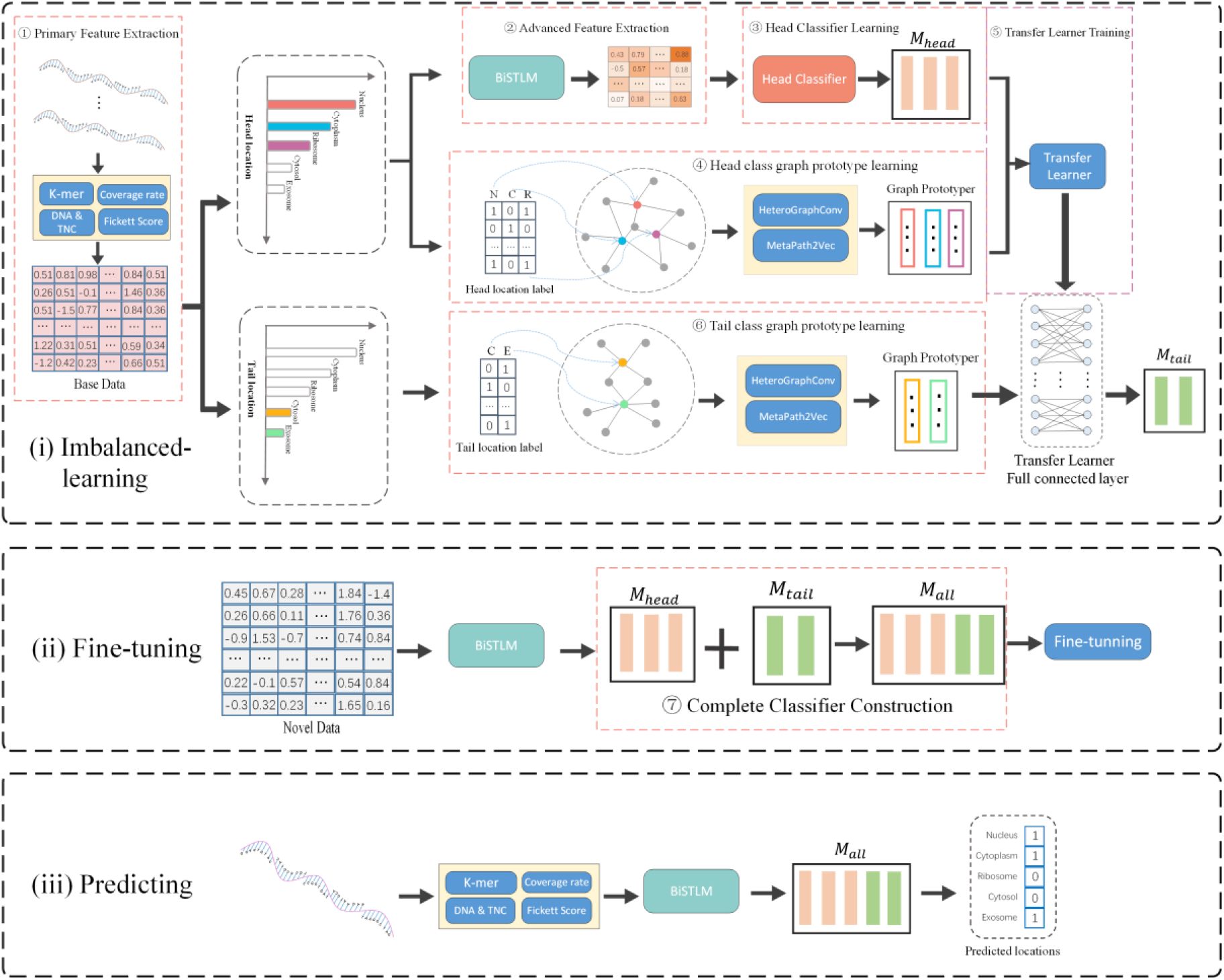
The overall architecture of GP-HTNLoc comprises three main components: (i) imbalanced learning based on graph prototypes, (ii) fine-tuning, and (iii) prediction. In the unbalanced learning phase, this study introduces a graph prototype module that obtains prototypical representations of labels from head label samples, which in turn trains a transfer learner to transfer rich categorization knowledge from the head class to the sample-scarce tail class (illustrated in the figure using the lncRNA dataset).

### 2.3 GP-HTNLoc

#### 2.3.1 Deep Feature Extraction

The deep feature extraction part contains primary and advanced feature extraction, the primary features are obtained from the original ncRNA sequences by four sequence feature extraction methods, and the advanced features are obtained by inputting the primary features into the BiLSTM with attention mechanism.

This study employs seven nucleotide sequence vectorization methods, namely k-mer nucleotide composition, Nucleic Acid Composition (NAC), Di-Nucleotide Composition (DNC) and Tri-Nucleotide Composition (TNC), regional coverage and nucleotide content differences, as well as the Fickett Score. These methods are utilized to extract primary features from the original RNA sequences. K-mer, NAC, DNC, and TNC are commonly used methods for extracting nucleotide sequence features. In this study, the extraction of these three features follows the same computational methods as described by Zhou et al. [17].Additionally, it has been shown that some short open reading frames (ORFs) of ncRNAs have the potential to encode micropeptides [29], and the 3’-UTR (3’ untranslated region) and 5’-UTR (5’ untranslated region) in non-ORF regions are equally biologically important as two important segments of RNA [30]. Therefore, follow features were included in the feature set: the overall coverage of the open reading frame region, the coverage of the 3’-UTR and the 5’-UTR, and the difference in guanine and cytosine content between the 3’-UTR and the 5’-UTR. Fickett and colleagues proposed that codons may exhibit asymmetric biases and nucleotide content, which can be utilized to distinguish between non-coding and protein-coding regions of a sequence [31]. Based on this observation, they introduced the Fickett Score to characterize the distinctiveness in nucleotide content and position. The exact calculation of the Fickett Score refers to wang et al [32].

After obtaining the primary features 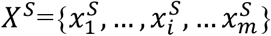 of the ncRNA sequence through the above multiple sequence feature extraction methods, we use a BiLSTM with attention mechanism [23] to further extract the deeper features of the sequence 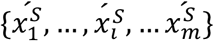,

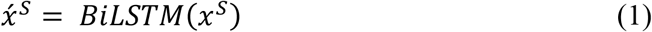

#### 2.3.2 Head Classifier Learning

In order to circumvent excessively complex classifier models that fail to yield robust generalization performance on datasets with limited samples, this study exclusively employs a multilayer perceptron devoid of bias terms as the classifier. The classifier solely consists of the weight matrix 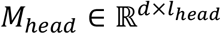, where *l*_*head*_ represents the number of head labels. The head classifier is trained utilizing the deep features extracted from sequences in *D*_*head*_, denoted as 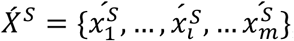,

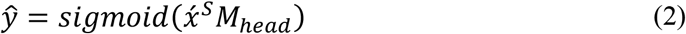

The weights of the head classifier, denoted as *M*_*head*_ are learned by minimizing the cross-entropy loss function. After multiple epochs of training on *D*_*head*_, we save the parameters of *M*_*head*_ that yield the best performance as the final head classifier parameters. The head classifier is trained on *D*_*head*_, which contains a substantial number of ncRNA samples, resulting in its superior classification capability. In the subsequent steps of the workflow, we aim to transfer this enhanced classification capability of the head classifier to the tail classifier, using graph prototype module and transfer learning technique.

#### 2.3.3 Graph Prototype Module

Graph representation learning is an important branch in the field of machine learning that aims to represent nodes and edges in graph data, such as social networks, recommendation systems, and bioinformatics, as vectors or embeddings. This helps in performing downstream tasks such as node classification, link prediction, community detection and recommendation [34]. Researchers have developed various graph embedding techniques to convert original graph data into high-dimensional vectors. Examples include graph convolutional networks (GCNs) [35], variational graph autoencoders (VGAEs) [36], graph attention networks (GATs) [37], and random walk-based models such as DeepWalk [38] and Node2Vec [39]. For general homomorphic graphs these methods show excellent graph embedding capabilities, but for heteromorphic graphs the above methods are not up to the task. Considering the complexity of heterogeneous graphs, researchers have developed representation learning methods specialized for heterogeneous graphs, such as Heterogeneous Graph Convolutional Networks (HGCN), MetaPath2Vec, etc., and these methods have achieved excellent performances in heterogeneous graph embedding in different scenarios [40].

In this study, we innovatively propose a graph prototype module, which utilizes the association information between ncRNA labels and samples to construct a heterogeneous graph, learns a high-dimensional embedding of labeled nodes on the heterogeneous graph using HGCN and MetaPath2Vec, and uses this high-dimensional embedding as a category prototype for subsequent model training. Figure 3 illustrates the workflow of the graph prototype module, taking the head class of the LncRNA dataset as an example. The process is delineated into the following steps:

**Figure 3.**
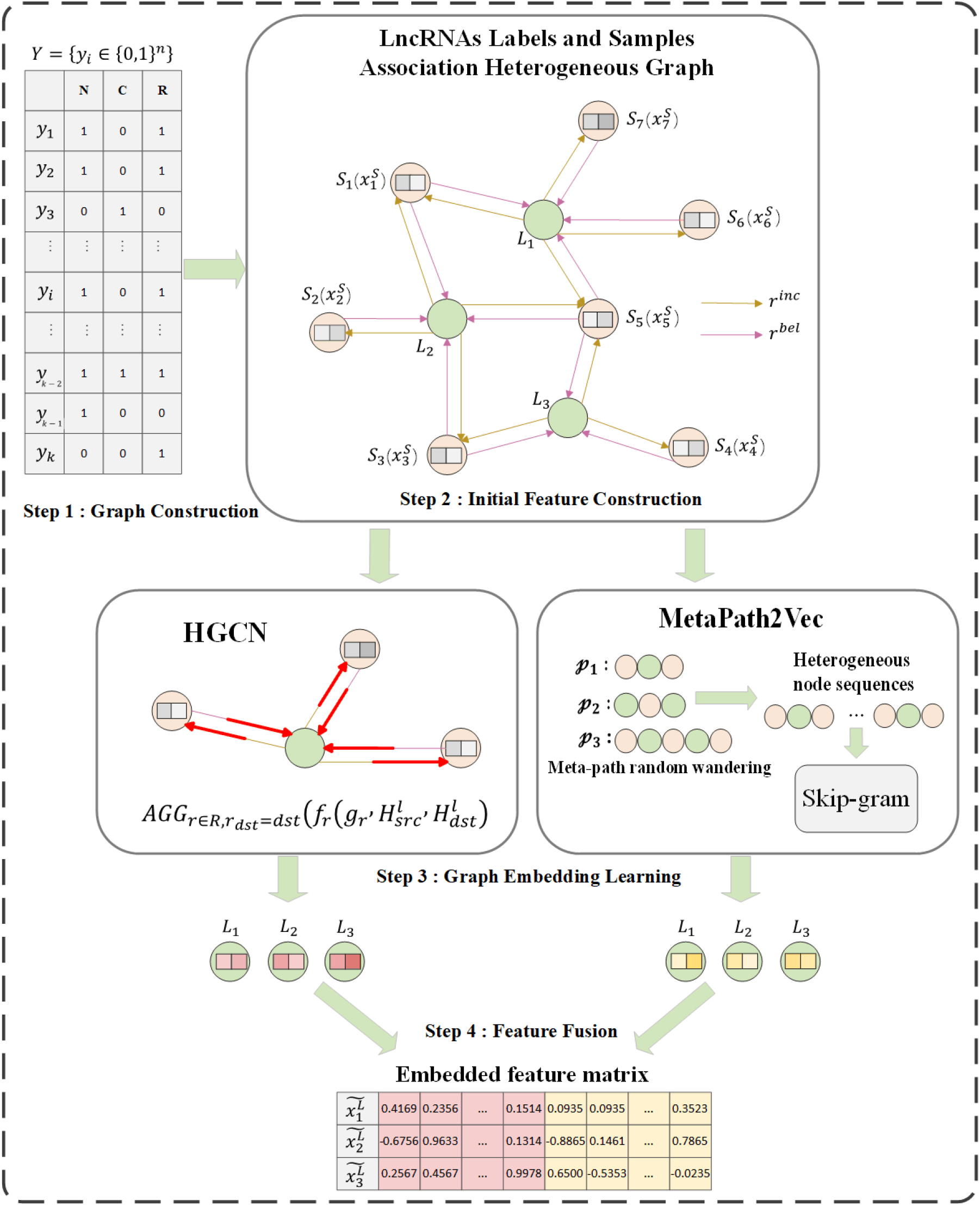
Workflow diagram of the graphical prototype module. Firstly, the labels and samples association heterogeneous graph is constructed from the label matrix, then the sample nodes are initialized with deep sequence features and the label nodes are initialized with the standard normal distribution. Following this, the label node embeddings are learned on the heterogeneous graph by using HGCN and MetaPath2Vec, respectively. Finally, the two types of embeddings are fused to obtain the final label embeddings as labeling prototypes.

##### Step 1 -- Graph Construction

A two-dimensional table can be generated from the localization label set *Y* = {y_*i*_ ∈ {0,1}^*n*^}, as depicted in Figure 3. In the table, rows represent different LncRNA samples, and columns represent different localization labels. If the position at the *i*-th row and *j*-th column is 0, it indicates an association between the LncRNA sample 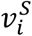 and the label 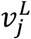. In this case, a directed edge is established from 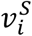 to 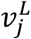 to signify that the sample belongs to the label, denoted as 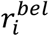. Similarly, a directed edge is established from 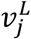 to 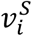 to represent the relationship indicating that the label includes the sample, denoted as 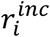. If a position in the label table is 0, it implies the absence of an association between the corresponding row and column samples and localization labels. In this scenario, no edge relationship is established between the corresponding label and sample nodes. The final result is the heterogeneous graph *G* representing the association between LncRNA labels and samples, denoted as *G* = (*V, R, X*), where *V* ∈{*V*^*S*^, *V*^*L*^ }. *V*^*S*^ represents LncRNA sample nodes (depicted as orange circles in Figure 3), 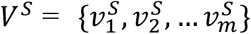, where *m* is the total number of LncRNA samples. *V*^*L*^ represents localization label nodes (depicted as green circles in Figure 3), 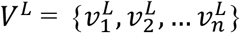, where *n* is the total number of LncRNA localization label nodes. *R* represents relationships between the two types of nodes, 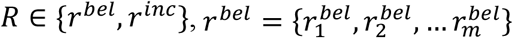, and 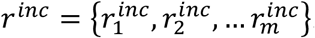. In Figure 3, *r*^*bel*^ is indicated by brown arrows, and *r*^*inc*^ is represented by purple arrows.

##### Step 2 -- Initial Feature Construction

*X* ∈ {*X*^*S*^, *X*^*L*^}, where *X*^*S*^ represents the features corresponding to all LncRNA sample nodes, 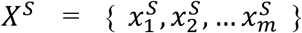. *X*^*L*^ represents the features corresponding to all label nodes, 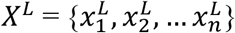. Initially, low-level features are extracted from the original sequences of each LncRNA sample to form *X*^*S*^. As for the label features *X*^*L*^, they are initialized using a standard normal distribution.

##### Step 3 -- Graph Embedding Learning

The embedding learning for label nodes is conducted on the heterogeneous graph *G* using both HGCN and MetaPath2Vec methods. The HGCN part performs graph convolution operations[41] on different relations *R* ∈ {*r*^*bel*^, *r*^*inc*^} of the heterogeneous graph separately, if multiple relations share the same target node type, their results are aggregated using the specified method, and if the relational graph has no edges, the module is not invoked, the formal definition is shown in equation 3. Eventually, after multiple rounds of propagation the sample feature information and graph structure information are aggregated on the labeled nodes to get the label embedding specific to HGCN.

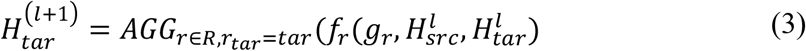

In this context, 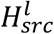 represents the activation matrix on the source nodes for the relationship *r*, and 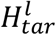 represents the activation matrix on the target nodes for the same relationship *r. g*_*r*_ denotes the subgraph on the heterogeneous graph *G* that exclusively contains the relationship *r*. The convolution result on relationship *r* is obtained through the function *f*_*r*_(·). Subsequently, the convolution results across multiple relationships are aggregated using the aggregation function *AGG*(·), producing the activation matrix 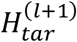 on the target nodes for the (*l* + 1)-th layer.

In MetaPath2Vec, the model utilizes meta paths to guide the process of random walks, generating heterogeneous node sequences. These node sequences are then inputted into Skip-Gram[42], enabling vectorization of the sequences and producing embeddings for the label nodes. In our task, the meta path “SLS” indicates that a certain LncRNA sample (S) belongs to a specific localization label (L), which in turn includes another (or the same) sample (S). Furthermore, meta paths are often used symmetrically, which helps in recursive guidance during random walks [27,43]. In this study, on the heterogeneous graph that associates ncRNA samples with labels, “SLS”, “SLSLS”, “LSL” and “LSLSL” and various other meta paths to guide the random tour of graph embedding learning. The experimental results show that when the meta path “SLSLS” is used, the performance is slightly better, and there is no significant difference in the final model performance of the other meta paths.

The labeled embedding 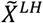, which pools the features of the samples, was obtained from HGCN, and the labeled embedding 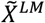, which is enriched with information about the graph structure, was obtained from MetaPath2Vec, and is formally described as follows:

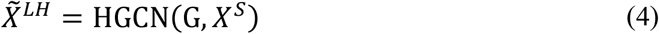

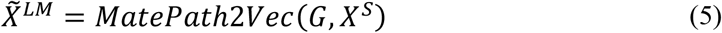

##### Step 4 -- Feature Fusion

To comprehensively leverage both sample feature information and graph structural details, this study horizontally concatenates the aforementioned two types of label embeddings to serve as the embedding for each label node. The formal description is as follows:

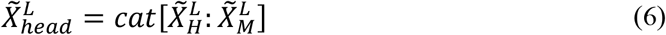

With the above four steps, we obtain labeled embeddings that contain rich information. In this study, 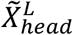 is used as a prototype of the head class for subsequent training of the GP-HTNLoc model.

#### 2.3.4 Transfer Learner Training

Up to this point, our study has trained the parameters *M*_*head*_ for the head classifier using sequence deep features on the relatively abundant samples in *D*_*head*_. Additionally, the graph prototype module has been utilized to construct prototypes for the head class.

To learn the mapping relationship from class prototypes to classifier parameters, we have established a transfer learner on the head class. The transfer learner consists of a multilayer perceptron without bias terms. The learnable weight matrix *W*_*transfer*_ ∈ ℝ^*d*×*d*^ serves as the trainable parameters for the transfer learner The training of the transfer learner parameters *W*_*transfer*_ involves minimizing the following loss function ℒ_*t*_:

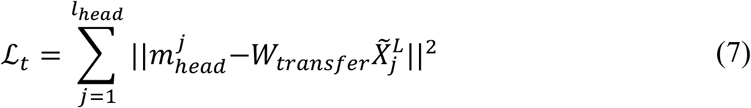

Where 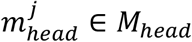 represents the parameters of the head class classifier, and 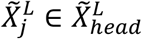 denotes the prototype for the head class.

#### 2.3.5 Complete Classifier Construction

On *D*_*tail*_, the graph prototype module is similarly employed to obtain prototype representations 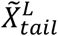 corresponding to the tail labels. Subsequently, the mapping from the tail class prototypes to the parameters of the tail classifier is accomplished using the previously trained transfer learner,

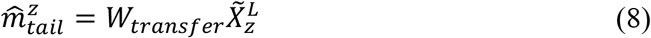

The prototype representation 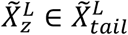 corresponds to the tail class, where 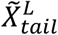 represents the prototypes for the tail class. Additionally, 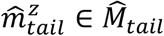 denotes the parameters of the tail class classifier. The construction of the final classifier parameters is outlined as follows:

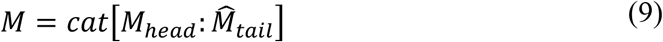

So far, we have resolved the imbalance in the head and tail categories of the ncRNA dataset and initially constructed a subcellular multilabel localization predictor.

#### 2.3.6 Fine-tuning and Prediction

The complete classifier *M*, obtained by directly concatenating *M*_*head*_ and 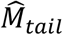, is likely to exhibit excellent performance solely on samples involving either head or tail class labels. However, due to the lack of optimization on data containing both head and tail class labels simultaneously, it may not achieve the anticipated performance on samples simultaneously involving both label types. In response, this study designed a fine-tuning module to minimally train the complete classifier parameter *M* using tail class data, called Novel Data, containing both head class labels and tail class labels. In specific experimental trials, we observed that merely one epoch of fine-tuning sufficed to yield significant improvements in *M* on data containing both head and tail class labels. Surprisingly, excessive fine-tuning reintroduces class imbalances that degrade model performance. So, in all subsequent code experiments, the fine-tuning phase was consistently restricted to a single epoch.

During the prediction phase, for a given novel ncRNA sequence, after extracting primary features through multiple sequence feature extraction methods, these primary features are fed into a BiLSTM network equipped with an attention mechanism to obtain advanced features 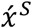. The corresponding label set 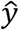 for this ncRNA sequence can then be obtained through the following process:

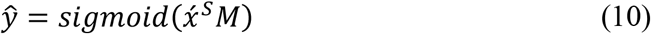

Where *M* represents the parameters of the complete multi-label classifier after fine-tuning. Finally, we obtained predictions of subcellular localization of ncRNA sequences by GP-HTNLoc.

### 2.4 Evaluation Metrics

Distinguished from typical binary and multiclass problems, multi-label classification entails unique evaluation metrics [44]. In this study, six commonly employed metrics in multi-label classification problems [16,17,18] are utilized to assess the performance of GP-HTNLoc. The six evaluative metrics include Average Precision (*AP*), Hamming Loss(*L*_*h*_), One-Error(*E*_*noe*_), Coverage (*Cov*), Accuracy (*ACC*), and Ranking Loss(*L*_*r*_). For the aforementioned evaluation metrics, higher values in Average Precision and Accuracy signify superior model performance, while smaller values in Hamming Loss, One-Error, Coverage, and Ranking Loss indicate better model performance.

### 2.5 Implementation details

GP-HTNLoc is implemented based on PyTorch[45], with the HeteroGraphConv(HGCN) and MetaPath2Vec modules in the graph prototype module implemented using DGL[46]. Both the head class classifier and fine-tuning utilize BCELoss for training, while the transformation learner is trained using MSELoss. We employed the Adam optimizer with a learning rate of 0.00001, along with default beta1 and beta2 values for model training.

## 3 Results

This study conducts rich code experiments on six benchmark datasets, including feature combination, parameter analysis, comparison experiments, case study, and ablation study, to fully demonstrate the excellent performance of GP-HTNLoc.

### 3.1 Comparison with different features

In order to identify the optimal combination of features, this study conducted experiments on three datasets, snoRNA, lncRNA and miRNA datasets, as mentioned in the previous sections. The experiments focused on all feature extraction methods described in ‘Section 2.3.1’. The extraction of these features was performed using Python code from the original RNA sequences, and the dimensions as well as the identifiers of each feature are shown in Table 1.

**Table 1.** Feature Dimensions and identifiers.

| Features | Dimensions | identifiers |
| --- | --- | --- |
| 2-mer | 16 | A |
| 3-mer | 64 | B |
| 4-mer | 256 | Z |
| Coverage-Rate | 5 | C |
| NAC | 4 | N |
| DNC | 16 | D |
| TNC | 64 | T |
| Fickett Score | 8 | F |

In this study, we inputted the aforementioned features, including their combinations, into the model and assessed the performance through 10-fold cross-validation. Due to the considerable number of feature combinations, we initially evaluated the performance of each feature independently and prioritized the top-ranked features for subsequent combinations. Table 2 presents a subset of superior-performing features and their combinations, detailing their performance across the three RNA datasets. Observing the table, it is evident that on the lncRNA dataset, the feature combination ABCDTF (2-mer, 3-mer, Coverage-Rate, DNC, TNC, Fickett Score) demonstrates the highest performance, achieving an *ACC* of 0.473 and an *AP* of 0.738. On the snoRNA dataset, feature T (TNC) attains the best performance with an *ACC* of 0.505 and an *AP* of 0.775. In addition, feature N (NAC) on the miRNA dataset achieves the highest performance with an *ACC* of 0.523 and an *AP* 0.771. For subsequent experiments, this study will employ the best feature or combination of features from each dataset.

**Table 2:** Performance of Different Feature Combinations on ncRNA Datasets.

| Dataset | Metrics | AB | N | D | T | Z | ABC | CDT | ABDTF | ABCDTF |
| --- | --- | --- | --- | --- | --- | --- | --- | --- | --- | --- |
| lncRNA | ACC | 0.403 | 0.369 | 0.395 | 0.422 | 0.455 | 0.425 | 0.438 | 0.465 | **0.473** |
|  | AP | 0.701 | 0.650 | 0.698 | 0.702 | 0.712 | 0.709 | 0.717 | 0.721 | **0.738** |
| | $E_{one}$ | 0.447 | 0.456 | 0.445 | 0.430 | 0.421 | 0.431 | 0.432 | 0.399 | **0.392** |
| | $L_r$ | 0.207 | 0.232 | 0.228 | 0.214 | 0.202 | 0.209 | 0.207 | 0.197 | **0.189** |
| snoRNA | ACC | 0.480 | 0.487 | 0.499 | **0.505** | 0.472 | 0.473 | 0.502 | 0.482 | 0.460 |
|  | AP | 0.745 | 0.758 | 0.767 | **0.775** | 0.760 | 0.731 | 0.761 | 0.723 | 0.702 |
| | $E_{one}$ | 0.253 | 0.249 | 0.245 | **0.239** | 0.275 | 0.262 | 0.243 | 0.249 | 0.275 |
| | $L_r$ | 0.223 | 0.231 | **0.200** | **0.200** | 0.221 | 0.226 | 0.208 | 0.230 | 0.243 |
| miRNA | ACC | 0.509 | **0.523** | 0.488 | 0.466 | 0.506 | 0.478 | 0.483 | 0.453 | 0.428 |
|  | AP | 0.763 | **0.771** | 0.733 | 0.703 | 0.752 | 0.717 | 0.714 | 0.699 | 0.682 |
| | $E_{one}$ | 0.318 | **0.301** | 0.323 | 0.365 | 0.315 | 0.328 | 0.322 | 0.393 | 0.402 |
| | $L_r$ | 0.183 | **0.176** | 0.219 | 0.227 | 0.188 | 0.231 | 0.217 | 0.245 | 0.269 |

### 3.2 Comparison with different parameters

#### 3.2.1 Different head and tail class divisions

In GP-HTNLoc, a training strategy that separates head and tail categories is employed to address the issue of significant imbalances in the number of samples for ncRNA subcellular localization. However, the degree of class imbalance varies from one dataset to another, and it is not possible to give uniform criteria for class division. To address this challenge, this study explores the performance of GP-HTNLoc under various head-tail class division scenarios for six datasets. The specific scenarios are outlined in Table 3. In Table 3, the “divisions” column represents the strategies for dividing head and tail categories. For instance, “2+3” indicates that, after arranging categories (subcellular localization labels) in descending order based on sample quantities, the top two categories with the highest sample counts are designated as head categories, while the bottom three categories with fewer samples are considered tail categories. Other division methods follow a similar principle.

**Table 3:**
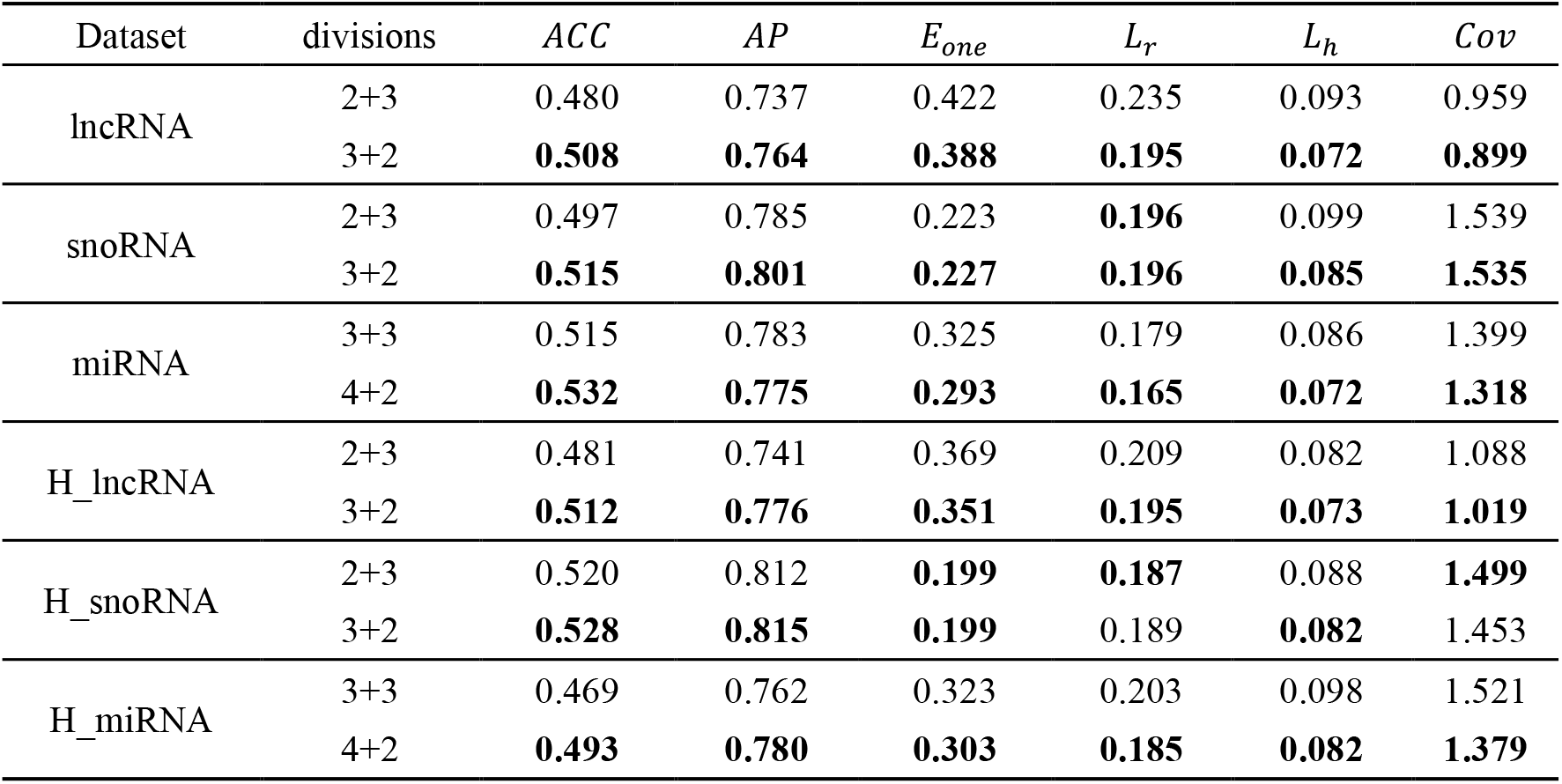
Performance of Different Head-Tail Class Division Strategies.

| Dataset | divisions | $ACC$ | $AP$ | $E_{one}$ | $L_r$ | $L_h$ | $Cov$ |
| --- | --- | --- | --- | --- | --- | --- | --- |
| lncRNA | 2+3 | 0.480 | 0.737 | 0.422 | 0.235 | 0.093 | 0.959 |
|  | 3+2 | **0.508** | **0.764** | **0.388** | **0.195** | **0.072** | **0.899** |
| snoRNA | 2+3 | 0.497 | 0.785 | 0.223 | **0.196** | 0.099 | 1.539 |
|  | 3+2 | **0.515** | **0.801** | **0.227** | **0.196** | **0.085** | **1.535** |
| miRNA | 3+3 | 0.515 | 0.783 | 0.325 | 0.179 | 0.086 | 1.399 |
|  | 4+2 | **0.532** | **0.775** | **0.293** | **0.165** | **0.072** | **1.318** |
| H_lncRNA | 2+3 | 0.481 | 0.741 | 0.369 | 0.209 | 0.082 | 1.088 |
|  | 3+2 | **0.512** | **0.776** | **0.351** | **0.195** | **0.073** | **1.019** |
| H_snoRNA | 2+3 | 0.520 | 0.812 | **0.199** | **0.187** | 0.088 | **1.499** |
|  | 3+2 | **0.528** | **0.815** | **0.199** | 0.189 | **0.082** | 1.453 |
| H_miRNA | 3+3 | 0.469 | 0.762 | 0.323 | 0.203 | 0.098 | 1.521 |
|  | 4+2 | **0.493** | **0.780** | **0.303** | **0.185** | **0.082** | **1.379** |

Table 3 demonstrates that, on the lncRNA and H_lncRNA datasets, the performance achieved by dividing three categories as head classes far surpasses the performance when two categories are designated as head classes. In both miRNA and H_miRNA datasets, the division with four head and tow tail classes showed better performance compared to the division with three head and three tail classes. Concerning the snoRNA and H_snoRNA datasets, the performance of the division with three head classes and two tail classes excels in *ACC, AP*, and *L*_*h*_ compared to the division of two head classes and three tail classes. However, the performance in *E*_*one*_, *L*_*r*_, and *Cov* between these two division strategies remains comparable, possibly attributed to a reduced sample size in these datasets compared to the other four. Overall, the trend suggests that GP-HTNLoc achieves better predictive performance when a larger number of categories are designated as head classes. This could be attributed to the tendency for head classifiers to learn more comprehensive classification information when faced with a greater variety of categories within the head classes. Consequently, this facilitates a more effective utilization of transferred meta-knowledge from the transfer learner onto the tail classes, thereby enhancing the overall model performance.

#### 3.2.2 Different graph prototype dimensions

In the graph prototype module of the GP-HTNLoc, class prototypes are obtained by concatenating embeddings learned from the graph using two methods: HGCN and MetaPath2Vec. The dimensions of the embeddings obtained by each method determine the dimensions of the prototypes, and these prototype dimensions are crucial for GP-HTNLoc. To investigate the impact of different prototype dimensions on the performance of GP-HTNLoc, this study conducted experiments on three datasets: lncRNA, snoRNA, and miRNA. The specific details are presented in Table 4 and Figure 4, where “Hgc-d” represents the embedding dimensions obtained through HGCN, “Mat-d” represents the embedding dimensions obtained through MetaPath2Vec, and “Total-d” represents the dimensions of the prototypes obtained by directly concatenating the aforementioned two embeddings.

**Table 4.**
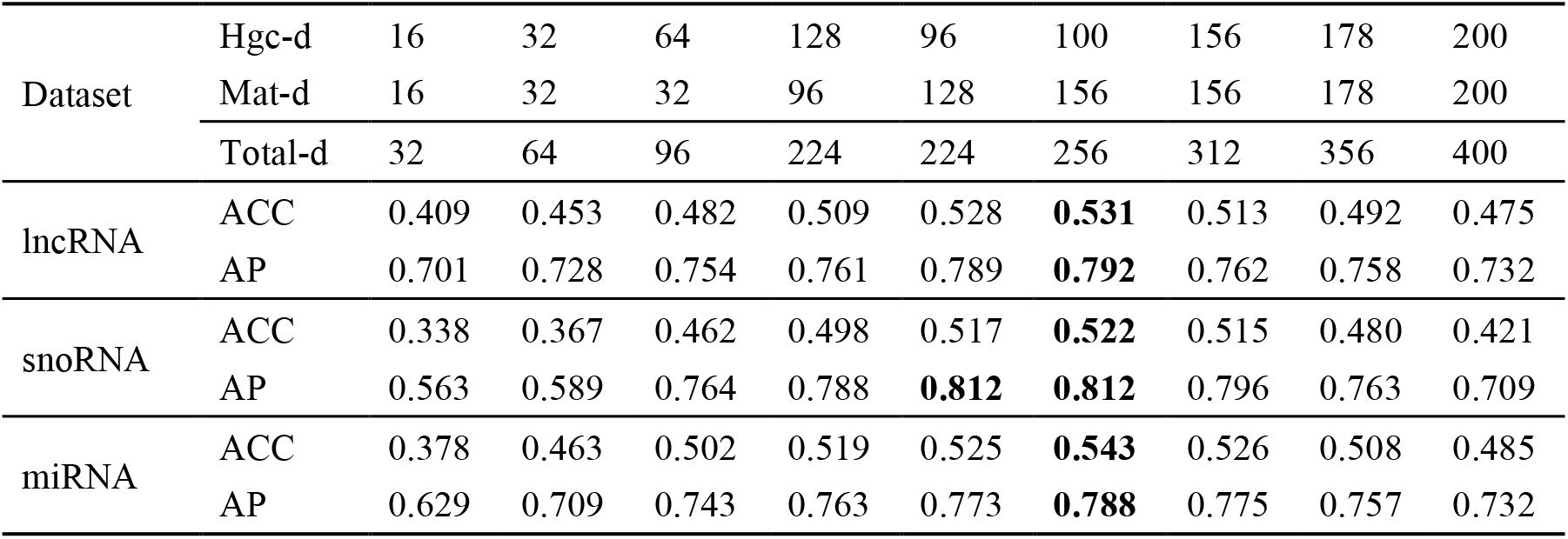
Performance of different embedding dimensions of the graph prototype module on three datasets.

**Figure 4.**
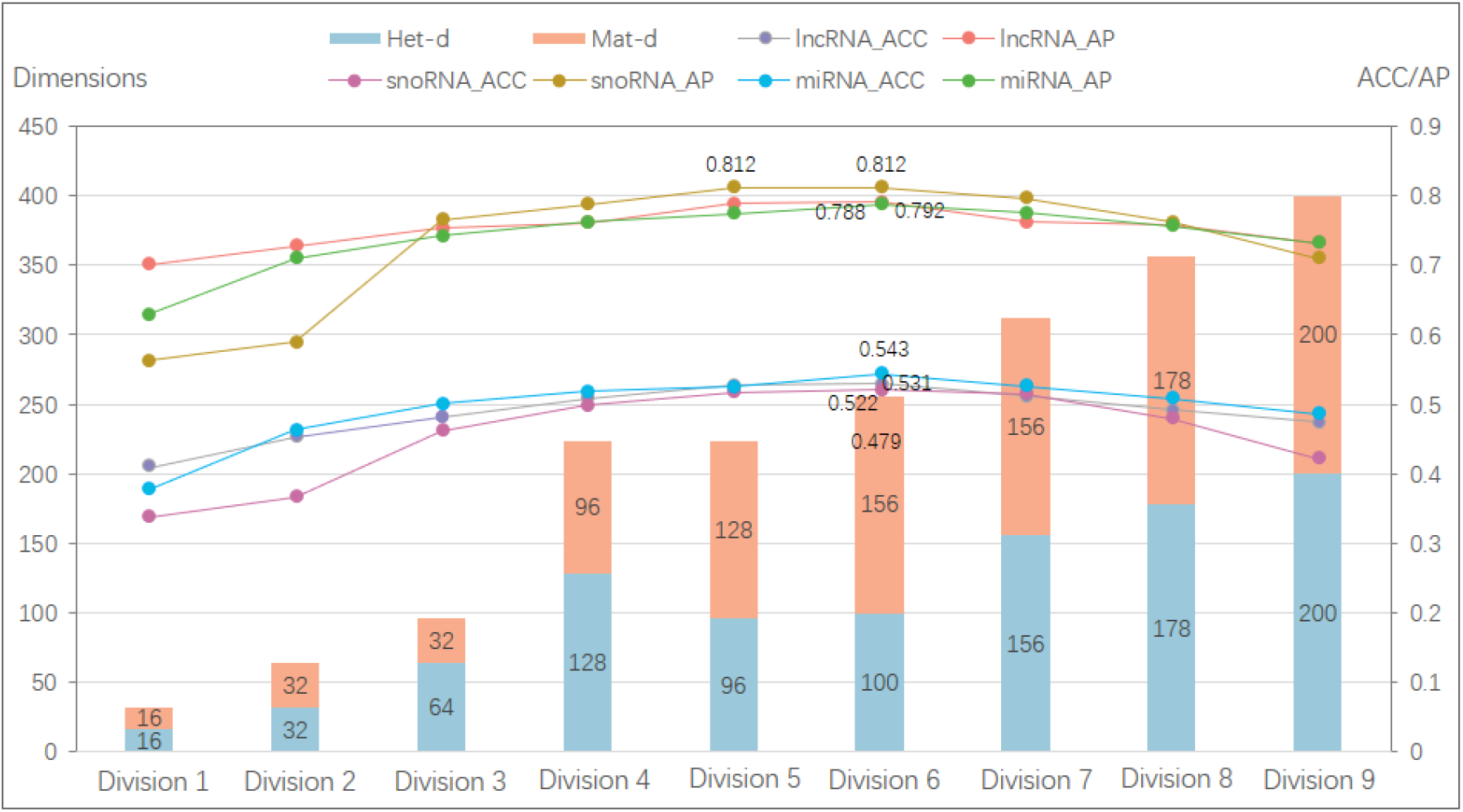
The performance of the prototype module with different embedding dimensions across three datasets

From Figure 4, it can be observed that initially, the model’s performance on each dataset improves with an increase in the dimensionality of graph embeddings. When the MetaPath2Vec embedding dimension is 156, the HGCN embedding dimension is 100, and the total prototype dimension is 256, the model consistently achieves the best performance across the three datasets. Subsequently, as the dimensions of the two embedding components continue to increase, the model’s performance gradually declines. In summary, GP-HTNLoc exhibits optimal performance when the graph prototype dimension is 256, as evidenced by the comprehensive evaluation across the three datasets.

### 3.3 Performance Analysis

Considering the impact of the graph prototype learning quality on the final model performance, we explored different numbers of training epochs for HGCN before performing performance evaluation. The experimental results show that HGCN reaches its optimal performance with 3 to 4 training epochs, while further increase in the number of training epochs leads to performance degradation instead. This may be due to the fact that 3 to 4 epochs are sufficient to aggregate all the sample features and graph structure information to the label nodes, while more training instead introduces noise.

To demonstrate the performance of GP-HTNLoc, this study employed optimal head-tail class partitioning, optimal graph embedding dimensions, and optimal heterogeneous graph embedding training epochs on each dataset. The performance of GP-HTNLoc was assessed through 10-fold cross-validation. The study computed all evaluation metrics over 10 runs of 10-fold cross-validation on all six datasets, along with the corresponding mean value, confidence interval, and standard deviation (significance level = 0.05). The detailed results are presented in Table 5. Additionally, we plot the model’s box-plot on all six indicators, as shown in Figure 5. The tables and figures illustrate that, GP-HTNLoc obtained small variance and deviation on six datasets, which proves its stable performance.

**Table 5.** Confidence means and variances of GP-HTNLoc for six metrics on each dataset.

| Datasets | ACC |  | AP |  | *E<sub>noe</sub>* |  |
| --- | --- | --- | --- | --- | --- | --- |
|  | mean | std | mean | std | mean | std |
| lncRNA | 0.528±0.0036 | 0.0058 | 0.792±0.0020 | 0.0033 | 0.367±0.0033 | 0.0054 |
| snoRNA | 0.522±0.0060 | 0.0096 | 0.812±0.0030 | 0.0049 | 0.219±0.0048 | 0.0077 |
| miRNA | 0.545±0.0036 | 0.0058 | 0.788±0.0015 | 0.0025 | 0.291±0.0020 | 0.0033 |
|  | *Cov* |  | *L<sub>r</sub>* |  | *L<sub>h</sub>* |  |
|  | mean | std | mean | std | mean | std |
| lncRNA | 0.882±0.0039 | 0.0063 | 0.167±0.0012 | 0.0020 | 0.062±0.0004 | 0.0006 |
| snoRNA | 1.526±0.0214 | 0.0345 | 0.188±0.0044 | 0.0071 | 0.076±0.0013 | 0.0021 |
| miRNA | 1.311±0.0091 | 0.0147 | 0.163±0.0018 | 0.0029 | 0.068±0.0007 | 0.0011 |
| Datasets | ACC |  | AP |  | *E<sub>noe</sub>* |  |
|  | mean | std | mean | std | mean | std |
| H_lncRNA | 0.527±0.0022 | 0.0036 | 0.785±0.0015 | 0.0025 | 0.342±0.0024 | 0.0039 |
| H_snoRNA | 0.532±0.0035 | 0.0057 | 0.821±0.0038 | 0.0062 | 0.193±0.0037 | 0.0059 |
| H_miRNA | 0.513±0.0050 | 0.0081 | 0.795±0.0030 | 0.0048 | 0.281±0.0050 | 0.0080 |
|  | *Cov* |  | *L<sub>r</sub>* |  | *L<sub>h</sub>* |  |
|  | mean | std | mean | std | mean | std |
| H_lncRNA | 0.998±0.0053 | 0.0086 | 0.183±0.0017 | 0.0028 | 0.067±0.0004 | 0.0006 |
| H_snoRNA | 1.486±0.0177 | 0.0285 | 0.180±0.0039 | 0.0063 | 0.075±0.0007 | 0.0012 |
| H_miRNA | 1.291±0.0101 | 0.0163 | 0.160±0.0018 | 0.0029 | 0.079±0.0009 | 0.0015 |

**Figure 5.**
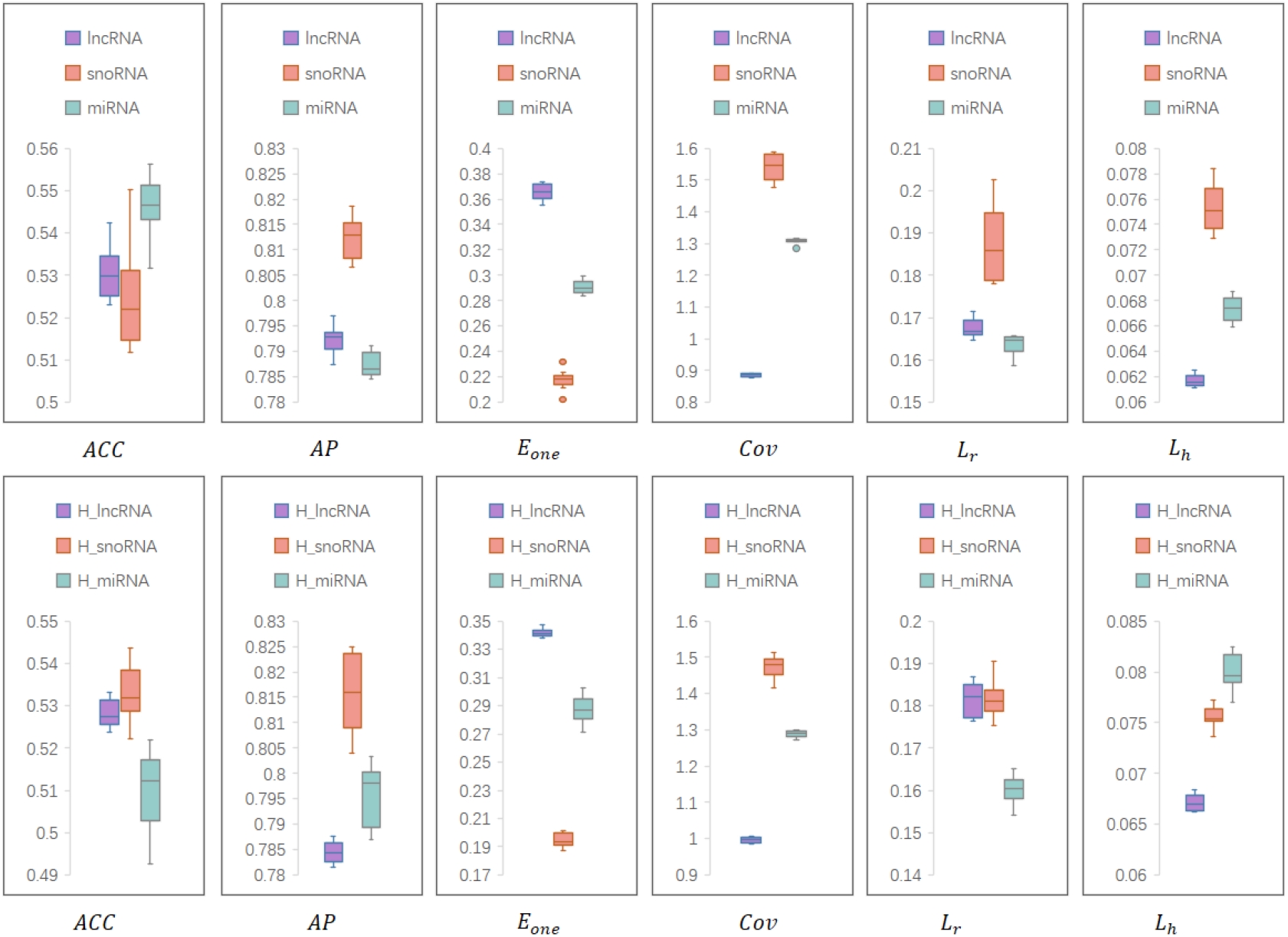
Boxplots of GP-HTNLoc on the RNA dataset and the human RNA dataset. Means, confidence intervals, and standard deviations (significance level=0.05) were calculated for the six evaluation metrics on all datasets.

To further validate the outstanding performance of GP-HTNLoc, this study conducted a comparative analysis against state-of-the-art models in the subcellular multi-label RNA localization domain. The comparative models included MKSVM proposed by Wang et al. [18] and MKGHkNN proposed by Zhou et al. [19]. The results are shown in Table 6, it is evident that GP-HTNLoc achieves an *ACC* of 0.528 on the lncRNA dataset, surpassing the current state-of-the-art model by 9.4%. Moreover, its average *AP* reaches 0.792, indicating a 3.3% improvement over the leading model. For the H_lncRNA dataset, GP-HTNLoc reached 0.527 on ACC, outperforming the best existing model by 9.3%, and 0.785 on AP, outperforming the existing model by 2.6%. Furthermore, GP-HTNLoc exhibits superior performance across all metrics on the H_snoRNA and snoRNA dataset in comparison to the current state-of-the-art model. On the miRNA dataset GP-HTNLoc outperforms the state-of-the-art model on four metrics except *ACC* and *Cov*. On the H_miRNA dataset our model outperforms existing models on *AP, E*_*one*_, *Cov, L*_*r*_, and the gap between our model and the state-of-the-art model is very small on *ACC* and *L*_*h*_.

**Table 6.**
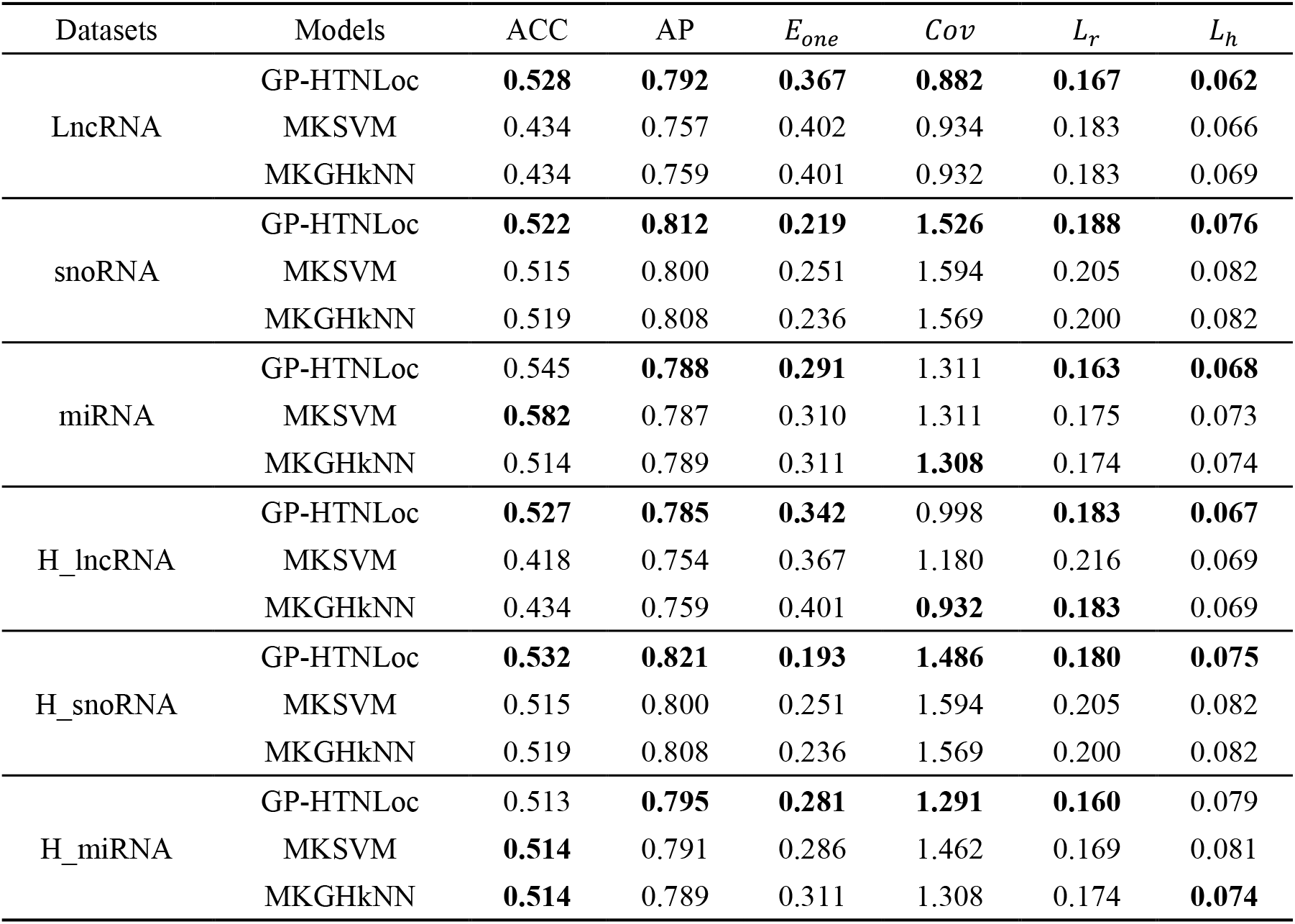
Comparison of GP-HTNLoc performance with state-of-the-art models.

This study further conducts a visual analysis of the model performance comparison, as illustrated in Figure 6, where the red bars represent the metrics corresponding to GP-HTNLoc. It is obvious from the figure that the GP-HTNLoc proposed in this study reaches the state-of-the-art in terms of subcellular localization of ncRNAs compared to existing advanced models.

**Figure 6.**
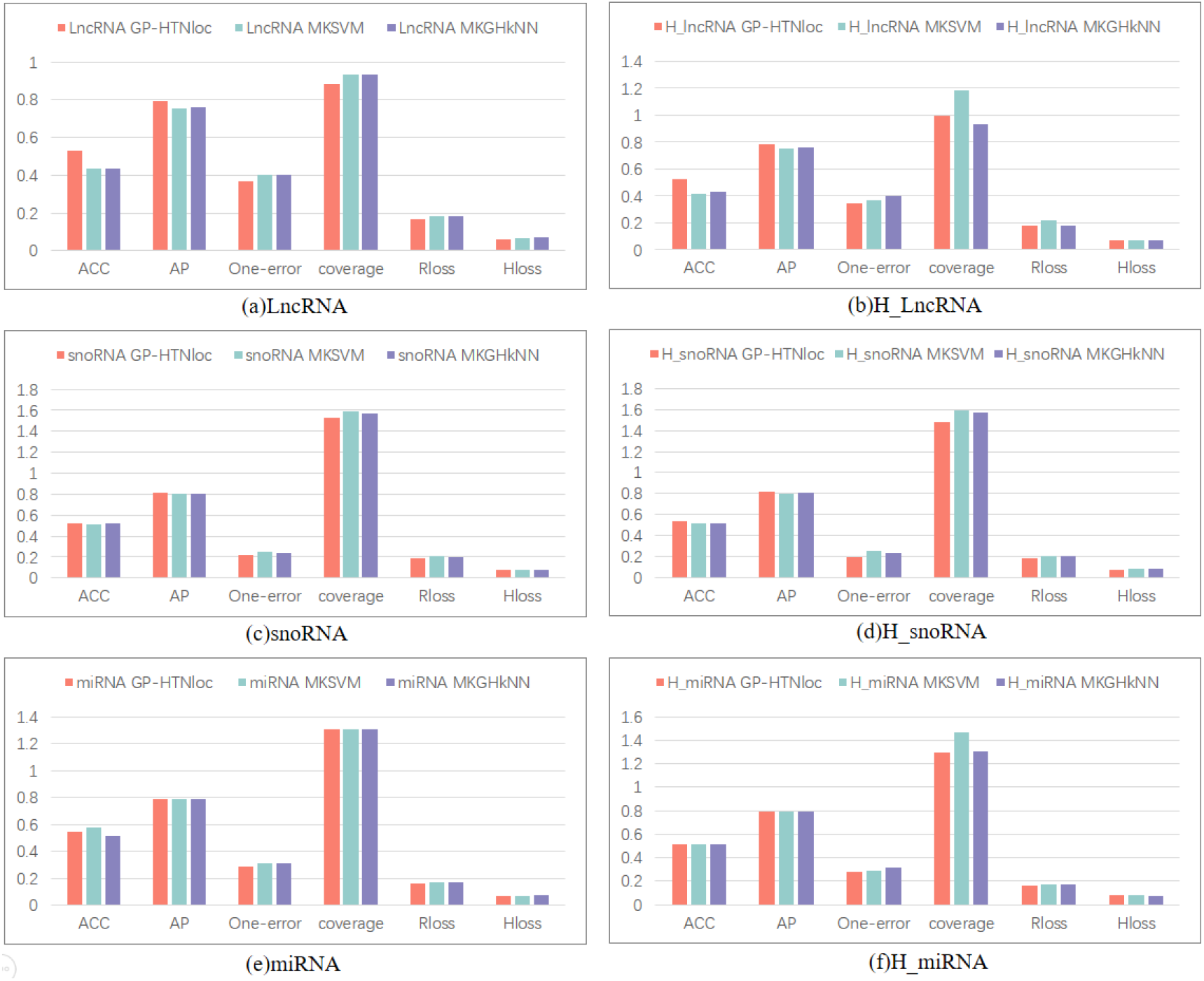
Performance comparison between GP-HTNLoc and SOTA models

### 3.4 Case Study

In order to further substantiate the reliability of GP-HTNLoc in practical scenarios of ncRNA multi-label localization prediction, we obtained CARL lncRNA (Cardiac Apoptosis-Related Long Non-coding RNA, NCBI:66774|Ensembl:ENSMUSG00000097638) from RNALocate v2[9]. The RNA Symbol for CARL is Carlr. It is noteworthy that this particular lncRNA is absent from the six training datasets utilized in our study. Research by Castellanos-Rubio et al. indicates that under basal conditions in mouse macrophages, Carlr predominantly exhibits nuclear localization, while after LPS stimulation, a majority of Carlr transcripts localize within the cytoplasm[47], suggesting the true subcellular localization of CARL lncRNA encompasses both the nucleus and cytoplasm. We conducted a case study using this lncRNA as an example to assess GP-HTNLoc. Employing the five-label classifier trained on the LncRNA dataset in Section 2.1, GP-HTNLoc predicted the multi-location subcellular localization of CARL lncRNA. The prediction process revealed sigmoid probability values for five subcellular locations—Nucleus, Cytoplasm, Ribosome, Cytosol, and Exosome—as follows: 0.553, 0.562, 0.173, 0.197, and 0.194, respectively. Using a threshold of 0.5 for class assignment, the final prediction resulted in Nucleus and Cytoplasm, aligning perfectly with the true subcellular locations of this lncRNA. Moreover, the considerable discrepancy in probability values between positive and negative classes in the sigmoid further underscores the discriminative capability of GP-HTNLoc.

### 3.5 Ablation study

To substantiate the pivotal role of the graph prototype module introduced in this study for enhancing the performance of GP-HTNLoc, we conducted ablation study on the six datasets used in the preceding sections. We tested GP-HTNLoc with the graph prototype module introduced in this study and a model using sample averaging as the prototype, referred to as AP-HTNLoc. As shown in Table 7, the results of the study indicate that replacing the graphical prototype module with the average prototype module resulted in varying degrees of deterioration of the model for all metrics across the six datasets. Consequently, these findings demonstrate that the introduced graph prototype module in this study effectively learns discriminative information from the complex sample label-associated heterogeneous graph, resulting in a substantial improvement in the predictive performance of the model. The ablation study underscore the significance of the graph prototype module as a critical component of GP-HTNLoc.

**Table 7.** Ablation study of graph prototype module.

| Datasets | Models | ACC | AP | $E_{one}$ | $Cov$ | $L_r$ | $L_h$ |
| --- | --- | --- | --- | --- | --- | --- | --- |
| LncRNA | GP-HTNLoc | **0.531** | **0.795** | **0.368** | **0.882** | **0.167** | **0.062** |
|  | AP-HTNLoc | 0.474 | 0.757 | 0.396 | 0.934 | 0.190 | 0.071 |
| snoRNA | GP-HTNLoc | **0.521** | **0.811** | **0.220** | **1.528** | **0.188** | **0.076** |
|  | AP-HTNLoc | 0.503 | 0.770 | 0.263 | 1.604 | 0.208 | 0.087 |
| miRNA | GP-HTNLoc | **0.539** | **0.782** | **0.293** | **1.315** | **0.166** | **0.069** |
|  | AP-HTNLoc | 0.515 | 0.737 | 0.323 | 1.341 | 0.179 | 0.078 |
| H_lncRNA | GP-HTNLoc | **0.528** | **0.786** | **0.342** | **0.997** | **0.182** | **0.067** |
|  | AP-HTNLoc | 0.468 | 0.755 | 0.382 | 1.119 | 0.207 | 0.079 |
| H_snoRNA | GP-HTNLoc | **0.530** | **0.820** | **0.193** | **1.488** | **0.180** | **0.075** |
|  | AP-HTNLoc | 0.515 | 0.800 | 0.243 | 1.598 | 0.205 | 0.083 |
| H_miRNA | GP-HTNLoc | **0.512** | **0.795** | **0.280** | **1.288** | **0.162** | **0.080** |
|  | AP-HTNLoc | 0.499 | 0.745 | 0.308 | 1.454 | 0.177 | 0.088 |

## 4 Discussion

The experimental results from the previous chapter reveal that the proposed ncRNA subcellular multi-label localization prediction model, GP-HTNLoc, exhibits significantly superior predictive performance on six benchmark datasets compared to existing state-of-the-art ncRNA subcellular localization prediction models. We analyze three key factors contributing to the success of GP-HTNLoc:

1. In contrast to traditional sequence feature extraction methods, we employ a two-stage feature extraction strategy. Initially, we combine various traditional feature extraction methods to extract primary feature representations from the ncRNA original sequences. Subsequently, we use a BiLSTM with attention mechanisms for deep sequence feature extraction. The primary features ensure the retention of as much original sequence information as possible. The attention mechanism in the BiLSTM focuses on the contextual information of the sequence while assigning different attention weights to different feature dimensions. Consequently, the derived high-level sequence features encompass profound discriminative information of ncRNA sequences, creating a more valuable feature space for the model’s subsequent graph prototype learning and final classification.
2. GP-HTNLoc introduces a learning paradigm of separate training for head and tail classes, effectively addressing the issue of class imbalance in ncRNA datasets without the need for resampling or loss weighting. This separate training approach not only avoids introducing noise but also preserves the original distribution characteristics of the dataset to a significant extent. The head and tail classifiers are trained using relatively balanced high-quality datasets, ensuring that the final integrated classifier performs well on class-imbalanced test datasets.
3. GP-HTNLoc introduces a novel graph prototype module that harnesses the association information between labels and samples for heterogeneous graph embedding learning. In this module, multiple ncRNA sample nodes in the heterogeneous graph may be associated with the same label node, and a single ncRNA sample node may also be concurrently associated with multiple label nodes. The implicit label association information embedded in this structure is deemed crucial for multi-label classification. Therefore, the graph prototype module overcomes the drawbacks of the One-vs-Rest strategy, which neglects label association information. Additionally, the learning of graph prototypes aggregates advanced features of sequences and extensively explores heterogeneous graph information through MetaPath2Vec. This process provides high-quality label prototypes for the training of the transfer learner.

Although GP-HTNLoc has achieved encouraging results, the current quantity of ncRNA samples used for GP-HTNLoc training remains limited. In future research, we aim to enhance model training by incorporating additional ncRNA sequences with reliable subcellular localization information from the latest ncRNA subcellular localization databases.

On another note, recent studies highlight the significance of cell line specificity in investigating subcellular environments. For instance, Lin et al. [48] noted that lncRNAs are typically expressed in a tissue-specific manner, and the subcellular localization of lncRNAs depends on the tissue or cell line in which they are expressed. Li et al. [49] pointed out that essential proteins are strongly influenced by the cellular environment, and essential proteins exhibit significant differences across cell lines. Therefore, in future research, we will incorporate cell line specificity into the model to further explore the prediction of subcellular multi-label localization for ncRNAs.

## 5 Conclusion

This study introduces a novel ncRNA subcellular multi-label localization model, GP-HTNLoc, which distinguishes itself from existing single-label localization models by simultaneously predicting multiple potential subcellular locations for each ncRNA sequence. Addressing the prevalent issue of class imbalance in RNA datasets, we adopt a training strategy that separates the more abundant head classes from the less numerous tail classes. In GP-HTNLoc, we innovatively introduce the graph prototype module, which requires only few samples to obtain label prototype representations containing rich graph structure information and potential label associations. This module enhances the model’s ability to predict subcellular multi-label localization for ncRNAs.

We conducted extensive experiments on GP-HTNLoc. Following the determination of optimal parameters for GP-HTNLoc, we compared its performance with state-of-the-art ncRNA subcellular multi-label localization models. The results revealed that GP-HTNLoc outperformed existing models on all datasets, achieving superior performance. Simultaneously, our conducted case studies highlight the robustness and applicability of GP-HTNLoc in practical scenarios of multi-label subcellular localization prediction for ncRNAs. Subsequently, we conducted ablation study on the model, and the results underscored the pivotal role played by the graph prototype module in enhancing the performance of GP-HTNLoc. In summary, GP-HTNLoc has demonstrated commendable performance in the field of ncRNA subcellular localization. We firmly believe that GP-HTNLoc will contribute significantly to advancing our biological understanding of ncRNA functions and mechanisms, further elucidating the regulatory patterns of ncRNA in the onset and progression of diseases.

## Data and code availability

The experimental codes and data sets for this study can be downloaded from https://github.com/han-skai/GP-HTNLoc.

## CRediT authorship contribution statement

Shuangkai Han (First Author): Model design; Method integration; Collection of the data; Most part of paper writing; Figure design; Visualisation; Experimental design; Most part of coding;

Lin Liu (Corresponding Author): Model design; Assistance in paper writing; Assistance in experimental design; Paper revision; idea Provision; Conceptualization, Funding Acquisition, Resources, Supervision, Writing - Review & Editing.

All authors have read and agreed to the published version of the manuscript.

## Acknowledge on funding

This work was supported by the Applied Basic Research Project in Yunnan Province (grant no. 202201AT070042), the project funding of the “Support Program of Xingdian Talents”, the National Natural Science Foundation of China (grant no. 61862067, U1902201), Yunnan Provincial Science and Technology Department-Yunnan University Double First-Class Joint Fund Key Projects (grant no. 2019FY003027) and National Key R&D Program of China (grant no. 2022YFC2602500).

